# Soluble RANKL exaggerates hindlimb suspension-induced osteopenia but not muscle protein balance

**DOI:** 10.1101/2020.09.03.281360

**Authors:** Toni L. Speacht, Charles H. Lang, Henry J. Donahue

**Affiliations:** Department of Cellular and Molecular Physiology, The Pennsylvania State University, College of Medicine, Hershey, PA 17033; Department of Orthopaedics and Rehabilitation, The Pennsylvania State University, College of Medicine, Hershey, PA 17033; Department of Biomedical Engineering, Virginia Commonwealth University, Richmond, VA

**Keywords:** Sarcopenia, Osteopenia, Bone-muscle interactions, Bone μCT, disuse, unloading

## Abstract

We examined the hypothesis that exaggerating unloading-induced bone loss using a combination of hindlimb suspension (HLS) and exogenous injections of receptor activator of nuclear factor kappa-B ligand (RANKL) also exaggerates muscle loss. Forty, male C57Bl/6J mice (16 weeks) were subjected to HLS or normal ambulation (ground control, GC) for 14 days. Mice received 3 intraperitoneal injections of either human recombinant soluble RANKL or PBS as control (n=10/group) at 24 hour intervals starting on Day 1 of HLS. GC + RANKL and HLS mice exhibited similar decreases in trabecular bone volume and density in both proximal tibias and distal femurs. However, RANKL affected trabecular number, separation, and connectivity density, while HLS decreased trabecular thickness. The combination of RANKL and HLS exacerbated these changes. Similarly, GC + RANKL and HLS mice saw comparable decreases in cortical bone volume, thickness, and strength in femur midshafts, and combination treatment exacerbated these changes. Plasma concentrations of P1NP were increased in both groups receiving RANKL, while CTX concentrations were unchanged. HLS decreased gastrocnemius weight and was associated with a reduction in global protein synthesis, and no change in proteasome activity. This change was correlated with a decrease in S6K1 and S6 phosphorylation, but no change in 4E-BP1 phosphorylation. Injection of RANKL did not alter muscle protein metabolism in GC or HLS mice. Our results suggest that injection of soluble RANKL exacerbates unloading-induced bone loss, but not unloading-induced muscle loss. This implies a temporal disconnect between muscle and bone loss in response to unloading.

## INTRODUCTION

Bone strength is reflective of both bone volume and bone quality. Bone volume (BV) is a dynamic balance between bone formation and bone resorption and BV decreases in unloading situations, such as prolonged bed rest, disuse due to aging, or in microgravity. For example, astronauts experience bone loss at a rate of 0.5% to 1.5% per month during spaceflight^1^ and femoral bone mineral density (BMD) had not fully recovered even a year after a 4 to 6 month stay on the International Space Station ^2^. Bone quality is determined by numerous variables including bone micro-architecture, microscopic damage, degree of mineralization, and bone turnover.

Previous findings suggest that muscle mass correlates with estimates of bone strength such as bone mineral density^3,4^. This relationship might be expected given that muscle is a primary source of mechanical stimuli for bone. Muscle atrophy as well as decreased contractile function also occur following prolonged bed rest ^5,6^, or spaceflight^7,8^.

The relationship of bone loss to muscle atrophy as a result of mechanical unloading is unclear. Previously, we showed that unloading-induced muscle atrophy precedes unloading-induced bone loss, which suggests that the loss of muscle may predispose, enhance, or accelerate bone loss^9^. However, we also reported that exaggerating unloading-induced muscle loss via a combination of HLS and casting does not exaggerate unloading-induced bone loss.^10^ In contrast, studies performed in both humans^11^ and animals^12^ suggest that maintenance of muscle mass during disuse mitigates bone loss. One theory is that the forces placed on bone from skeletal muscle are maintained, thereby partially conserving bone mass. Alternatively, muscle also secretes “myokines” that can influence bone metabolism^13^. To further explore the relationship between bone and muscle during unloading, we examined whether exacerbating unloading-induced bone loss would accelerate unloading-induced muscle loss.

Receptor activator of nuclear factor k-B ligand (RANKL) is a member of the tumor necrosis factor (TNF) superfamily, and is essential for osteoclast differentiation, activation, and survival^14,15^. RANKL is expressed by osteoblasts and binds to its receptor, RANK, on the surface of osteoclasts to stimulate differentiation and activation, leading to bone resorption. Osteoprotegerin (OPG) is a soluble decoy receptor to RANKL, thereby inhibiting bone resorption by preventing RANKL binding to RANK. The ratio of RANKL:OPG regulates bone resorption at the cellular level.

In this study, mice were suspended by their tail so that the hindlimbs were prevented from load-bearing. To exacerbate bone loss, soluble RANKL was injected into mice undergoing HLS. Studies have shown that exogenous administration of soluble RANKL in mice decreased bone volume, density, and strength similar to that produced by ovariectomy^16,17^. Lloyd et al injected 10-week-old female C57Bl/6J mice twice-daily with RANKL for 10 days, which resulted in an 84% decrease in trabecular bone volume fraction (BV/TV) and a 9% decrease in cortical volume ^16^. Others have induced significant bone loss by using only 3 injections of soluble RANKL to induce loss in 50 hours^17^. Additionally, OPG deficient (OPG -/-) mice have early onset osteoporosis characterized by a significant decrease in bone volume and strength^18,19^.

We found that HLS plus RANKL injection decreased trabecular and cortical bone volume compared to HLS alone. However, this loss did not exacerbate muscle atrophy or protein synthesis or degradation in mice experiencing both HLS and RANKL. These results demonstrate a temporal disconnect between muscle loss and bone loss, and suggest that maintaining muscle mass may not attenuate unloading-induced bone loss.

## MATERIALS & METHODS

### Animal procedures

Forty male C57BL/6J mice (Jackson Labs; Bar Harbor, ME) that were 16 weeks ± 3 days old at experimental day 0 were used. This age of mouse was used because it is skeletally mature and has been routinely used in spaceflight^20^ and ground-based simulation ^21^ studies. Mice were fed ad libitum standard rodent chow (2018 Teklad Global 18% Protein Rodent Diet; Envigo, Indianapolis, IN), maintained on a light/dark cycle of 12:12 hours, and housed at a constant temperature (25°C). One week before the start of experiments, mice were transferred to HLS cages to acclimate while ambulating normally. Mice were weighed and then randomly assigned to one of four groups (ground control (GC) + vehicle, GC + RANKL, HLS + vehicle, or HLS + RANKL; n=10/group). On day 1, mice received an intraperitoneal injection of either phosphate buffered saline (PBS; vehicle; Mediatech Inc., Manassas, VA) or human recombinant soluble RANKL (1 mg/kg; OYC Americas, Vista, CA). Mice received a total of 3 injections at 24 hour intervals. This concentration and time course was previously found to induce osteoporotic bone loss and decrease BMD for 4 weeks^17^. HLS was maintained for 14 days. At study endpoint, tibias, femurs, gastrocnemius, and quadriceps were harvested and blood samples were collected. Animal procedures were approved by the Penn State Institutional Animal Care and Use Committee (Protocol #46451).

### Hindlimb suspension

We used a modified version of the HLS model first described by Morey-Holton and Globus^22^ and previously described by our laboratory^10,23^. Mice were anesthetized with isoflurane (2% in O_2_; Phoenix Pharmaceuticals, Inc.; Burlingame, CA) and two strips of bandage tape braided around the tail, secured to a swivel fishhook attached to a string. The string was coiled around the cross bar at the top of the cage, such that the animal achieved an approximately 30° elevation until the hindlimbs were no longer in contact with the cage floor. Two mice were suspended per modified rat cage in this manner, but were placed at opposite ends to prevent physical contact. Standard corn cob bedding was placed beneath a wire mesh insert that allowed animals to access food and water ad libitum. GC mice were housed in a similar cage environment, but not suspended. Animals were inspected twice daily to assess general level of activity, responsiveness, and appearance. Each mouse received 1 ml of warmed (37 °C) sterile 0.9% saline (Baxter, Dearfield, IL) subcutaneously for resuscitation.

### Micro-computed tomography

Micro-computed tomography (microCT) was performed to assess micro-architectural properties of bone at baseline and study endpoint as previously described^10^. A Scanco vivaCT 40 (Scanco Medical AG; Brüttsellen, Switzerland) was used to scan the right tibia and femur from each animal. Cortical bone at the midshaft from the tibia and femur was assessed in 22 slices. Additionally, trabecular bone from the proximal tibia and distal femur (immediately distal and proximal to the epiphyseal plate, respectively) was assessed in 72 slices. Instrument settings and image reconstruction was performed exactly as previously described by our laboratory^10^. Cortical bone parameters reported herein include: total area inside the periosteal envelope (Tt.Ar), bone area (Ct.Ar), area fraction (Ct.Ar/Tt.Ar), thickness (Ct.Th), marrow area (Ma.Ar), tissue mineral density (TMD), cortical porosity (Ct.Po), and polar moment of inertia (pMOI). In addition, trabecular parameters were: total volume (TV), bone volume (BV), bone volume fraction (BV/TV), number (Tb.N), thickness (Tb.Th), separation (Tb.Sp), connectivity density (Conn.D), bone and mineral density (BMD). Above mentioned endpoints are reported as per guidelines^24^. Due to a hardware malfunction, we lost some baseline cortical scans resulting in a lower sample size (ranging from 4-10) for cortical microCT data.

### Biomechanical testing

Bones stored at −80 °C after MicroCT scanning were thawed and mechanically tested to failure via three-point bending using a Bose Electroforce (Bose; Eden Prairie, MN) and WinTest 4.0 software (TA Instruments; New Castle, DE) as previously described^9^. Flexural support spans were 8 mm. A loading rate of 1 mm/minute was applied in the anterior to posterior direction and a force (F) outcome measured. From a plot of F vs. displacement (d), bending rigidity (slope of the linear portion of the curve) and ultimate bending energy (area under the curve until the max force) were calculated.

### Serum markers of bone formation and resorption

At the study endpoint, 0.5mL of whole blood was collected into heparinized syringes from the vena cava of all mice. Blood was centrifuged for 10 minutes at 10,000 rpm at 4°C and plasma collected and frozen at −20°C until assayed. Type I collagen N-terminal propeptide (P1NP; Immunodiagnostic Systems Inc,; Gaithersburg, MD), C-terminal telopeptides (CTX; Immunodiagnostic Systems Inc), and osteoprotegerin (OPG; R&D Systems, Minneapolis, MN) were quantified with ELISA kits according to the manufacturer’s protocols and using a SpectraMax190 microplate reader (Molecular Devices, San Jose, CA) with SoftMax Pro software (Molecular Devices).

### In vivo protein synthesis

Protein synthesis in the gastrocnemius and quadriceps was determined in vivo on day 14 in all four groups of mice. Protein synthesis was determined in the freely fed state using the SUnSET method and an antibody against puromycin (Kerafast, Boston, MA)^25,26^. Conscious mice were injected intraperitoneal with puromycin (0.04 μmol/g body wt; 1 ml/100 g body wt) 30 minutes prior to tissue collection. Mice were anesthetized with isoflurane at 28 minutes and blood collected 2 minutes later. Thereafter, the gastrocnemius and quadriceps from each leg were rapidly excised. A portion of each muscle was freeze-clamped, stored at −80 °C, and processed as previously described^27^. Western blotting procedures were performed to visualize puromycin incorporation into muscle protein^26,28^. Furthermore, fresh muscle was homogenized and Western blotting was used for the determination of selected proteins; quantitative real-time (qRT)-PCR was performed on a separate piece of muscle, as described below.

### Western blot analysis

The phosphorylation state of 4E-BP1 and S6K1 has been typically used to assess the in vivo activation of mTORC1 (mechanistic target of rapamycin complex 1) which is a protein kinase that is central in regulating protein synthesis via various different inputs (e.g., growth factors, hormones, energy status, and mechanical stress)^29^. Thus, changes the phosphorylation of 4E-BP1 and S6K1 as well as S6 (which is phosphorylated by S6K1) is directly proportional to changes in muscle protein synthesis. Fresh muscle was homogenized (Kinematic Polytron; Brinkmann, Westbury, NY) in ice-cold homogenization buffer as previously described^10^. Western blot analysis was used to determine the relative amount of total and phosphorylated S65 (Bethyl Laboratories, Montgomery, TX) eukaryotic initiation factor 4E binding protein-1 (4E-BP1), and total and phosphorylated (T389; Cell Signaling Technology, Boston, MA) ribosomal protein S6 kinase (S6K) 1 and ribosomal S6 (S240/244; Cell Signaling Technology).

### Estimates of protein degradation

In vitro proteasome activity was determined in skeletal muscle as described previously^30^. To exclude nonproteasomal degradation, samples were assessed in the absence and presence of a proteasome inhibitor. Proteasome activity is reported as nmol·minute^−1^·mg protein^−1^.

### RNA extraction and real-time quantitative PCR

Muscle was homogenized in Tri-reagent (Molecular Research Center, Inc.; Cincinnati, OH) followed by chloroform extraction, as previously described^10^. Thereafter, the Qiagen RNAase mini kit (Qiagen; Valencia, CA) protocol was followed. RT-qPCR was performed for the genes encoding muscle RING-finger 1 (MuRF1) and atrogin-1 ^31^. Gene expression was presented in reference to controls using the comparative quantitation method 2^−ΔΔCt^ ^32^.

### Statistical analysis

Data are shown as mean ± standard error. A two-way ANOVA was used to evaluate differences between the four groups, with post-hoc Student-Neuman-Keuls test when the interaction was significant (p<0.05). GraphPad InStat was used for statistical analyses.

## RESULTS

### Tibia trabecular bone microarchitecture

HLS significantly affected trabecular bone architecture of proximal tibias as assessed by microCT. Vehicle-treated HLS mice lost 32% of BV/TV, compared to a 17% age-related decrease in GC mice (**Fig. 1A**). GC + RANKL mice demonstrated losses similar to those seen in HLS mice (−34%). Mice in the HLS + RANKL group lost significantly more BV/TV (−61%) than HLS alone. Mice exposed to HLS alone had a 17% decrease in Tb.Th compared to a 1% decrease in GC. However, injection of RANKL in GC or HLS mice had no effect on Tb.Th (−1% and −17%, respectively). Further, there was no difference in Tb.N, Conn.D, or Tb.Sp between GC and HLS mice (−6% vs −7%; −13% vs −4%; and +10% vs +10%, respectively). GC + RANKL mice lost 22% of Tb.N and 38% of Conn.D, and increased Tb.Sp by 36%, while the combination of HLS + RANKL had the most detrimental effects, a 31% decrease in Tb.N, 68% decrease in Conn.D, and 52% increase in Tb.Sp. HLS also decreased BMD (−22%) compared to GC (−9%), with a similar loss occurring in GC + RANKL mice (−22%). The greatest decrease in BMD was detected in HLS + RANKL mice (−43%). BV was decreased in HLS (−24%) compared to GC (−13%), with RANKL exacerbating the effect (−32% in GC and −57% in HLS). There was no difference in TV among any of the four groups.

**Figure 1.**
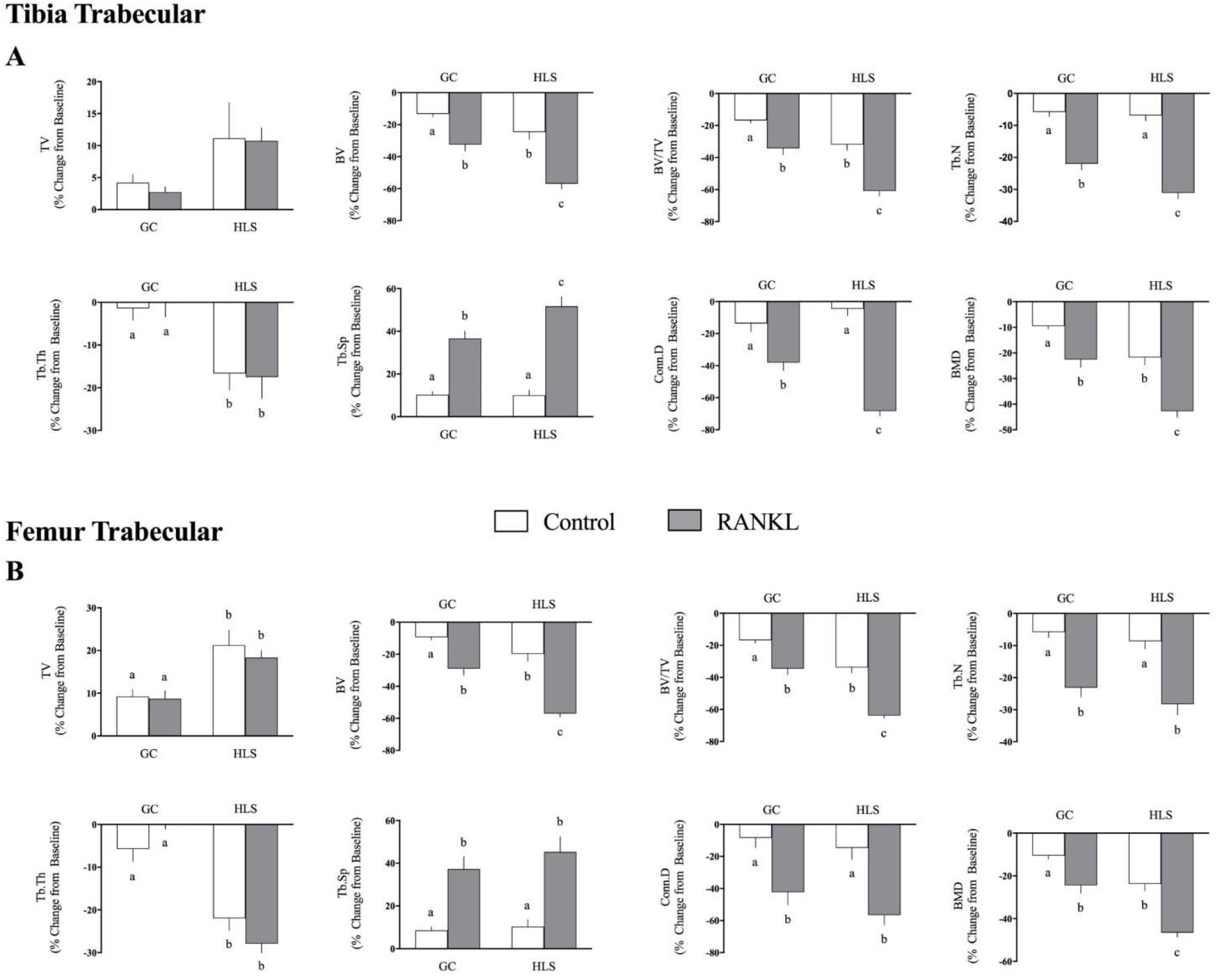
Effect of hindlimb suspension (HLS) ± RANKL on trabecular micro-architectural parameters obtained via microcomputed tomography scans of proximal tibia (A) and distal femur (B). Measured parameters include changes from baseline in total volume (TV), bone volume (BV), bone volume fraction (BV/TV), trabecular number (Tb.N), trabecular thickness (Tb.Th), trabecular separation (Tb.Sp), connectivity density (Conn.D), and bone mineral density (BMD). Values are means ± SEM; n = 10 for each group. Values with different letters are significantly (p < 0.05) different by 2-way ANOVA.

### Femur trabecular bone microarchitecture

HLS and RANKL also decreased trabecular bone of distal femurs (**Fig. 1B**). HLS decreased BV/TV by 34% compared to a 17% decrease in GC mice. A 34% decrease in BV/TV was also observed in GC + RANKL mice. Again, HLS + RANKL mice lost the most bone with a 64% decrease in BV/TV. HLS also decreased Tb.Th by 22%, with no further decrease in HLS + RANKL mice (−28%), compared to a 6% decrease in GC. HLS had no effect on Tb.N or Tb.Sp; however, RANKL injection in both GC and HLS mice produced a significant decrease. GC + RANKL mice lost 23% of Tb.Th and HLS + RANKL mice lost 28%, compared to the corresponding vehicle controls (−6% and −9%, respectively). Tb.Sp was increased 37% and 45% with RANKL injection in GC and HLS mice, compared to 8% and 10% increases in vehicle controls. Conn.D was decreased similarly in RANKL injected mice (−42% GC and −56% HLS), but significantly decreased compared to vehicle injected controls (−8% GC and −14% HLS). HLS and GC + RANKL had similar effects on trabecular BMD (−24% each) compared to GC (−10%). The loss of BMD was exacerbated in HLS + RANKL mice (−46%). BV decreased in both with HLS and GC + RANKL mice having similar losses (−19% and −29%, respectively), compared to GC (−9%). Again, HLS + RANKL exacerbated the loss (−57%). In femurs, TV was not affected by RANKL injection, but HLS increased TV by 21% (HLS) and 18% (HLS + RANLK) compared to GC (+9% each).

### Tibia cortical bone architecture

HLS alone had no effect on cortical bone of tibia midshafts for any cortical parameters measured (**Fig. 2A**). The injection of RANKL in HLS mice did not alter any of the measured parameters; however, in GC mice, RANKL decreased Ct.Ar (−8% vs +0.5%, +2%, and −1%), Ct.Ar/Tt.Ar (−9% vs −2%, −1%, and −2%), and Ct.Th (−15% vs −6%, −3%, and −5%). There was no difference among any groups for Tt.Ar, Ma.Ar, TMD, Ct.Po, or pMOI.

**Figure 2.**
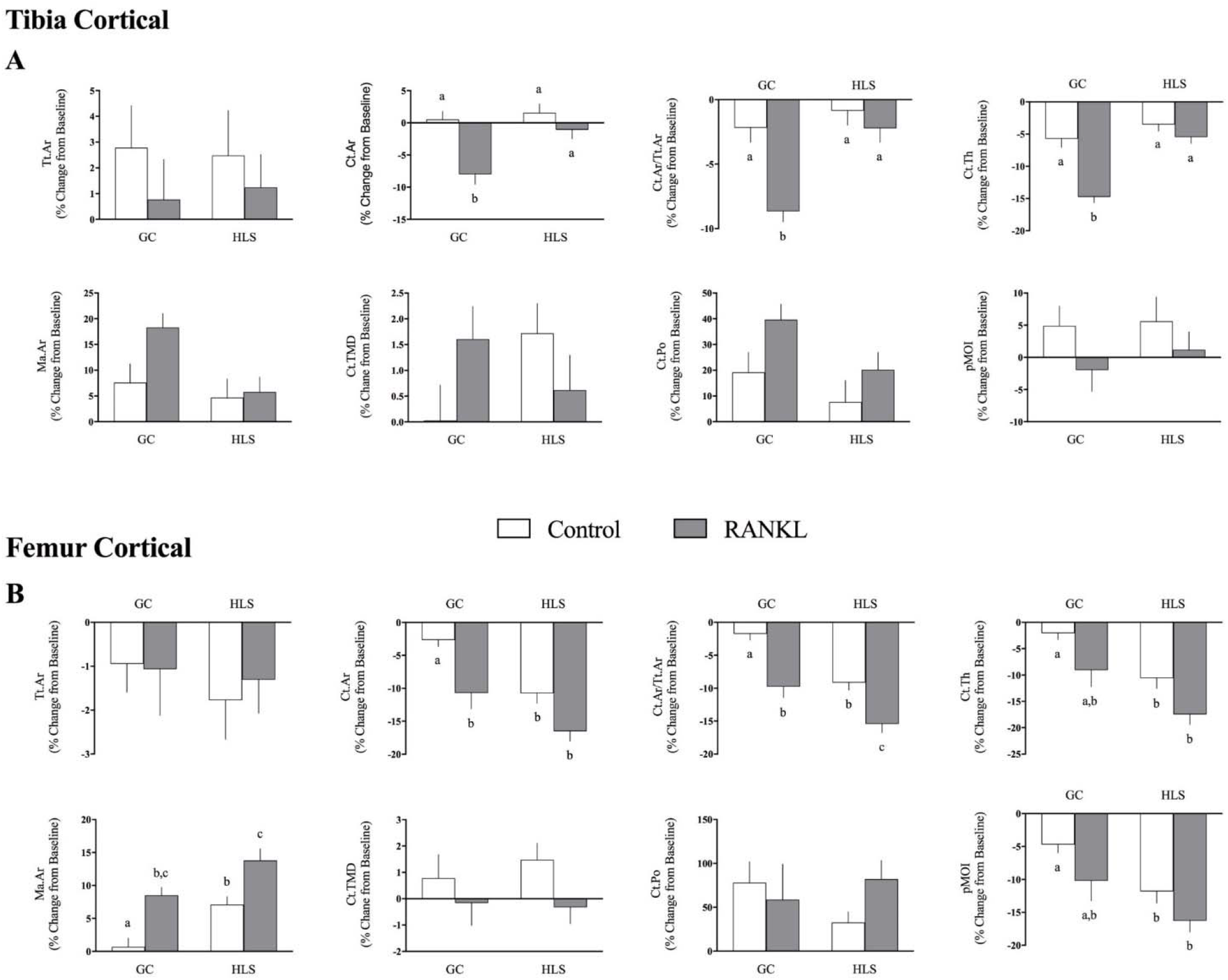
Effect of hindlimb suspension (HLS) ± RANKL on cortical micro-architectural parameters obtained via microcomputed tomography scans of tibia (A) and femur midshafts (B). Measured parameters include changes from baseline in total area (Tt.Ar), cortical area (Ct.Ar), cortical area fraction (Ct.Ar/Tt.Ar), cortical thickness (Ct.Th), marrow area (Ma.Ar), cortical tissue mineral density (Ct.TMD), cortical porosity (Ct.Po), and polar moment of inertia (pMOI). Values are means ± SEM; n = 4-10 for each group. Values with different letters are significantly (p < 0.05) different by 2-way ANOVA.

### Femur cortical bone architecture

Mice exposed to HLS lost 11% Ct.Ar, 9% Ct.Ar/Tt.Ar, 11% Ct.Th, and 12% pMOI in femur midshafts, compared to GC mice (−3%, −2%, −2%, and −5%, respectively; **Fig. 2B**). Similar results were observed in GC + RANKL mice (−11%, −10%, −9%, and −10%, respectively). HLS + RANKL mice only had further decreases in Ct.Ar/Tt.Ar (−15%), compared to HLS and similar decreases in Ct.Ar (−16%), Ct.Th (−17%), and pMOI (−16%). The Ma.Ar was increased comparably in both GC + RANKL (+8%) and HLS (+7%) mice compared to GC (+.7%). HLS + RANKL increased Ma.Ar (+14%) to a greater extent than HLS alone. There were no differences among any groups in Tt.Ar, TMD, or Ct.Po.

### Biomechanical Testing

HLS decreased stiffness compared to GC as determined by 3-point bending (Table 1). There was a comparable decrease in stiffness in GC + RANKL mice. Mice undergoing HLS + RANKL had even less stiffness than HLS mice and displayed a decreased peak force necessary to break bone compared to the GC groups. The ultimate bending energy did not differ between groups.

**Table 1.** Effect of hindlimb suspension (HLS) ± RANKL on biomechanical testing performed via three-point bending. Displacement (d) and force (F) were measured and used to calculate the ultimate bending energy and stiffness. Values are means ± SEM; n = 10 for each group. Values with different letters are significantly (p < 0.05) different by 2-way ANOVA.

|  | Ground Control |  | HLS |  |
| --- | --- | --- | --- | --- |
|  | Vehicle | RANKL | Vehicle | RANKL |
| **Stiffness (N/mm)** | 95.58 ± 3.41<sup>a</sup> | 86.32 ± 4.59<sup>a,b</sup> | 79.7 ± 4.47<sup>b</sup> | 67.5 ± 2.78<sup>c</sup> |
| **Peak Force (N)** | 16.37 ± 0.59<sup>a</sup> | 15.54 ± 0.52<sup>a</sup> | 14.62 ± 0.61<sup>a,b</sup> | 13.36 ± 0.46<sup>b</sup> |
| **Ultimate Bending Energy (Nmm)** | 2.53 ± 0.24 | 2.24 ± 0.16 | 2.3 ± 0.28 | 2.09 ± 0.11 |

### Serum markers of bone formation and resorption

The plasma concentration of P1NP was not different between GC and HLS groups without RANKL (**Fig. 3**). However, the injection of RANKL significantly increased P1NP levels in GC and HLS mice (60% and 73%, respectively), compared to their corresponding control. Plasma concentrations of CTX and OPG were not different among any groups at study endpoint.

**Figure 3.**
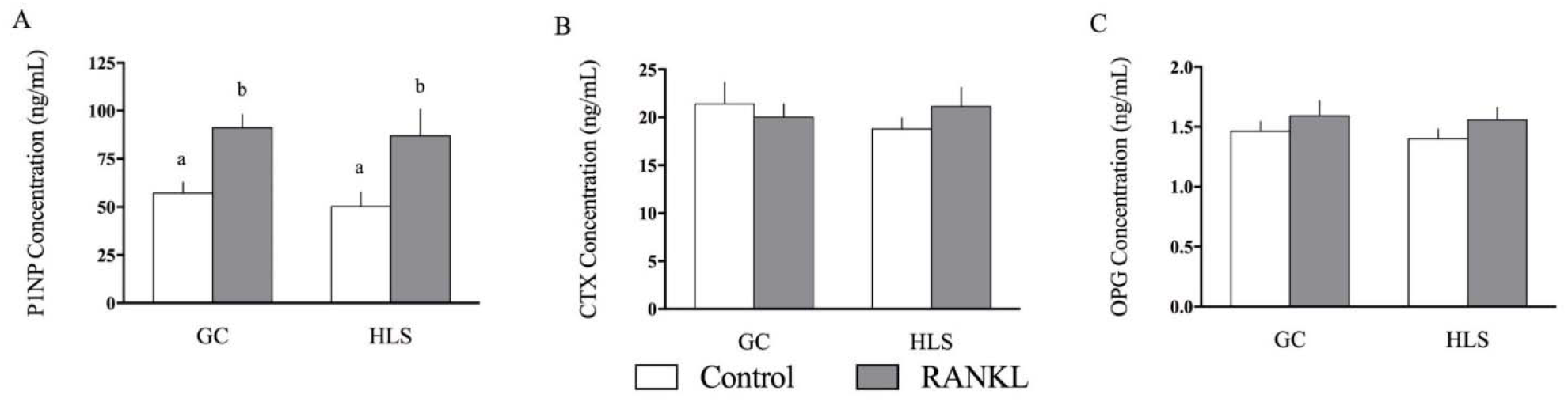
Plasma concentrations of markers of bone formation (type I collagen N-terminal propeptide; P1NP; A), bone degradation (C-terminal telopeptides; CTX; B), and osteoprotegerin (OPG; C) at study endpoint. Values are means ± SEM; n = 10 for each group. Values with different letters are significantly (p < 0.05) different by 2-way ANOVA.

### Muscle protein metabolism

Mice subjected to HLS demonstrated a decrease in gastrocnemius mass (−16% compared to GC) that was associated with a reduction in global protein synthesis (−29% vs. GC; **Fig. 4**). This latter change was associated with a reduction in the phosphorylation state of S6K1 and S6 (−41% and −32%, respectively), but not 4E-BP1. The injection of RANKL did not significantly alter muscle protein metabolism in GC mice nor did it alter the HLS-induced decrease in mass, protein synthesis or S6K1/S6 phosphorylation. Similar HLS-induced changes were detected in the quadriceps muscle, and again none of the above endpoints were altered by RANKL (data not shown). There was no difference in proteasome activity, or the mRNA content of MuRF1 or atrogin-1 among any groups in gastrocnemius (**Table 2**) or quadriceps (data not shown).

**Figure 4.**
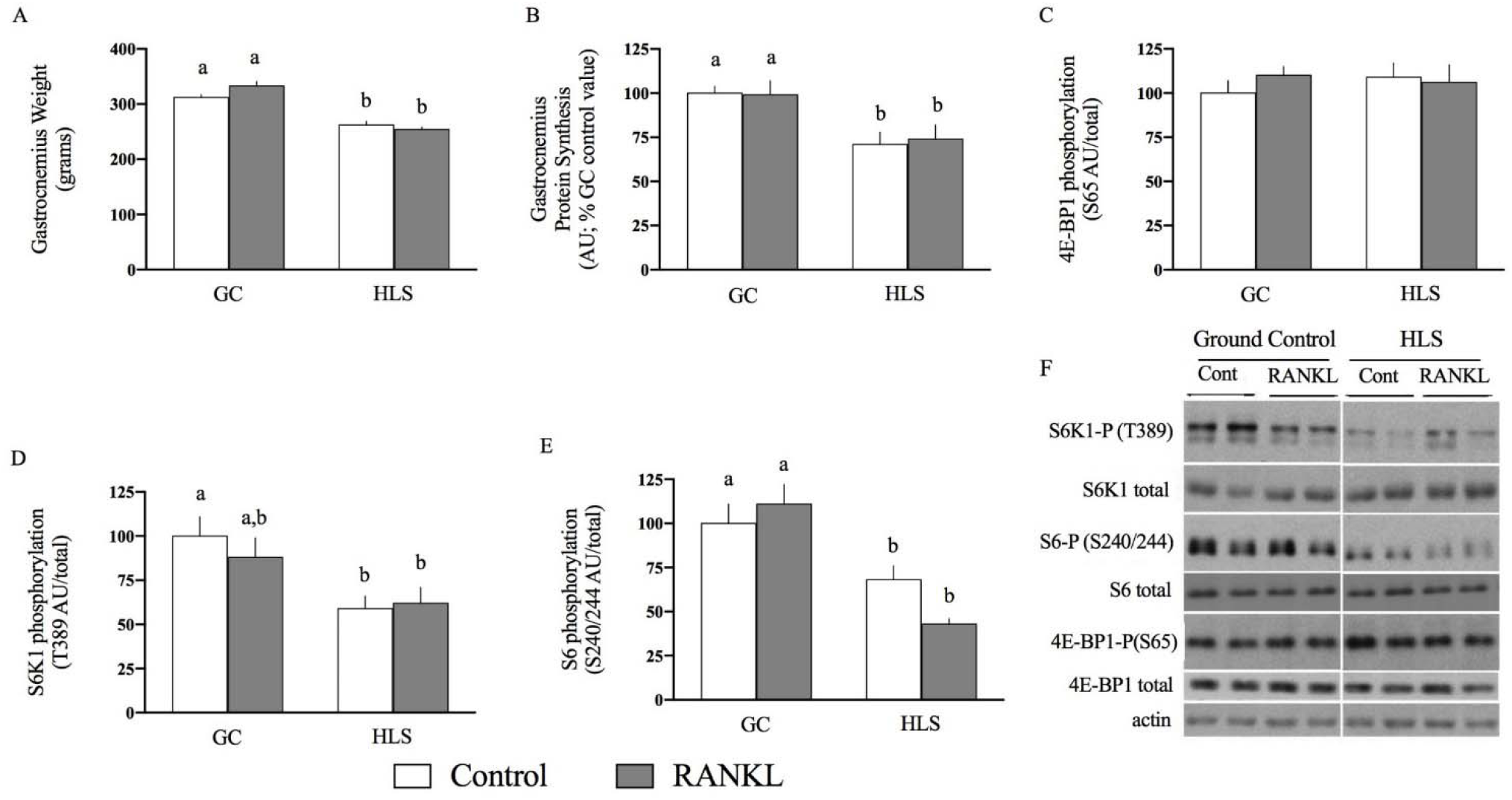
Effect of hindlimb suspension (HLS) ± RANKL on gastrocnemius weight (A), in vivo protein synthesis (B), 4E-BP1 phosphorylation (C), S6K1 phosporylation (D), and S6 phosphorylation (E). A representative Western blot is also presented (F). Values are means ± SEM; n=10 for each group. Values with different letters are significantly (p <0.05) different by 2-way ANOVA.

**Table 2.** Effect of hindlimb suspension (HLS) ± RANKL on proteasome activity, and mRNA levels of MuRF1 and Atrogin 1. Values are means ± SEM; n = 10 for each group. For each endpoint, there were no significant (p<0.05) differences between groups by 2-way ANOVA.

|  | **Ground Control** |  | **HLS** |  |
| --- | --- | --- | --- | --- |
|  | **Vehicle** | **RANKL** | **Vehicle** | **RANKL** |
| **Proteasome Activity**<br>*($\mu\text{mol}/\text{minute}/\text{mg protein}$)* | 2.5 ± 0.4 | 2.3 ± 0.3 | 3.1 ± 0.4 | 2.9 ± 0.6 |
| **MuRF1 mRNA**<br>*(AU/GAPDH)* | 1.00 ± 0.09 | 1.05 ± 0.12 | 1.07 ± 0.11 | 1.11 ± 0.12 |
| **Atrogin-1 mRNA**<br>*(AU/GAPDH)* | 1.00 ± 0.12 | 0.98 ± 0.08 | 1.13 ± 0.22 | 1.18 ± 0.19 |

## DISCUSSION

In thisstudy we used a novel approach to exacerbate bone loss without affecting muscle loss directly by injecting HLS mice with recombinant RANKL. We confirmed previous findings that hindlimb suspension alone^9,10,33^, as well as RANKL alone^16,17,34^, results in bone loss. We expanded on those findings and found that HLS plus RANKL exacerbated decreases in bone volume, density, and strength. We also unexpectedly found that mice receiving exogenous RANKL had increased markers of bone formation, suggesting that RANKL has a negative feedback loop regulating bone formation.

HLS and RANKL individually had similar effects on trabecular bone volume and density parameters, effects that were exacerbated when these catabolic stimuli were combined. However, HLS and RANKL affected different micro-architectural parameters measured by microCT. Our results suggest that RANKL induces bone resorption by decreasing the number of trabeculae as opposed to decreasing the thickness of each trabeculae. It is possible that osteoclasts positive for RANK receptors on trabecular bone are clustered together on trabeculae that would allow RANKL to increase osteoclast activity on certain trabeculae and not others. There is evidence that localization of RANK-expressing osteoclast precursors correlates with progression of inflammation in a collagen-induced arthritis model^35^ and to wound sites in periodontal defects in sheep^36^, therefore it is possible the RANK-expressing osteoclasts are localizing to specific trabeculae. This would allow for the decrease in trabecular number and connectivity density, while increasing trabecular separation, without affecting trabecular thickness. In regards to HLS, as the whole bone is experiencing uniform unloading, it is not surprising that trabecular thickness decreases equally across all trabeculae, and therefore the number of trabeculae does not change. The observation that these micro-architectural changes were exacerbated in the HLS + RANKL group in tibias of mice but not femurs, indicates a greater sensitivity to HLS + RANKL in tibia trabecular bone. The increase in total volume seen in distal femurs of HLS mice suggests periosteal bone formation may be occurring.

In femur midshafts, similar results were seen in most parameters of GC + RANKL and HLS mice, with the combination causing an exacerbated loss in bone. The increased marrow area with no change in total area suggests that HLS and RANKL both increase endocortical resorption, while turnover at the periosteal surface is unchanged. Cortical parameters in tibia midshafts were not affected by HLS, while RANKL administration in GC mice saw similar decreases as in femurs. The polar moment of inertia data from femurs correlate to our three-point bending data, signifying decreased bone strength in HLS mice and an even greater decreased strength in HLS + RANKL mice. These data suggest that the femur midshaft is more susceptible to changes from the combination of HLS and RANKL than tibia midshafts. This is opposite of what we found in the trabecular bone, but supports a previous study that reported whole femur BMD was more sensitive than whole tibias to the combination of HLS and ovariectomy^37^.

Plasma concentrations of P1NP, an indicator of bone formation, were not affected by HLS; however, they were increased in both GC and HLS mice that received RANKL injections. Plasma concentrations of CTX, which reflects bone resorption, were not different among any groups. These results suggest that bone loss in response to HLS and RANKL occurs rapidly and levels return to baseline thereafter. The increased P1NP levels in RANKL injected mice, but no other groups, suggest that increased RANKL plays a role in signaling for increased bone formation. This does not occur in HLS only mice, so this is not a compensatory mechanism due to bone resorption, but rather a signaling mechanism in response to RANKL. This supports previous theories that “reverse signaling” occurs with the TNF family members^38^. More recent studies have shown that low dose (0.125 mg/kg) administration of RANKL stimulates osteoblasts both in vitro and in vivo^39,40^. Buchwald and colleagues showed that RANKL was also able to induce regulatory T cells, thus engaging a negative feedback loop that stimulates osteoblasts^40^. While our P1NP results support these studies, it is important to note that we utilized a much higher dose of RANKL (1 mg/kg vs 0.125 mg/kg) than the other studies, so our results suggest that even high levels of RANKL can induce a negative feedback loop, but the net result will still be bone resorption.

Gastrocnemius weight was decreased in HLS mice, which corresponded to a decrease in global protein synthesis. Protein synthesis is primarily regulated by the mammalian target of rapamycin (mTOR) that phosphorylates S6K-1 and 4E-BP1, thereby stimulating translation. The decrease in muscle mass also corresponded to a decrease in S6K1 and S6 phosphorylation, but not 4E-BP1 phosphorylation. This difference in phosphorylation of downstream targets of mTOR is likely due to mTOR having a higher affinity to S6K1 and S6 than 4E-BP1^41^. These results confirm our previous work that showed muscle atrophy in response to HLS was mediated by a reduction in mTOR activity^9,10,23^. The injection of RANKL had no effect on muscle mass, protein synthesis, or mTOR activity.

We also assessed markers for muscle protein degradation. One mechanism that mediates protein degradation is ubiquitin-proteasome-dependent proteolysis^42–44^. Proteins targeted for degradation are tagged by ubiquitin by E3 ubiquitin ligases, specifically atrogin 1 and MuRF1 in striated muscle^45,46^. We found that neither HLS nor RANKL altered in vitro proteasome activity. Similarly, mRNA expression of atrogin 1 and MuRF1 were not altered in any groups, which agrees with our earlier findings that atrogin 1 and MuRF1 are increased at 7 days, but not 14 days after HLS^9,10^. These results suggest that high levels of exogenous RANKL (1mg/kg) do not affect protein balance and muscle mass. While RANK receptors are found in skeletal muscle^47^, it seems that RANKL has a more physiological impact on bone than muscle.

In this study, we were able to successfully exaggerate unloading-induced bone loss and examine the effects of this on muscle mass. Unexpectedly, we were able to exacerbate osteopenia without altering the resultant sarcopenia. Other studies from our lab have also suggested a disconnect between muscle and bone in response to unloading. Lloyd et al.^9^ reported that muscle loss precedes bone loss in response to unloading, suggesting a dominant role for muscle over bone in the homeostatic maintenance of these tissues. However, we also showed that casting plus HLS exacerbated muscle loss but did not exacerbate bone loss^10^. We proposed that this was due to either the relatively short duration of HLS (14 days) and that the bone required more time to manifest these changes or there was a level of temporal disconnect between the tissues. As we were able to induce greater bone loss in the same time period in the present study, such data would support the idea that muscle and bone are not as integrally connected in their response to unloading as originally believed. Krause et al.^23^ also found that the combination of high energy radiation (as experienced by astronauts in space) and HLS resulted in decreased bone volume and density, while quadriceps weight and protein synthesis were unchanged.

There have been multiple studies that show maintenance of muscle mass during disuse can attenuate bone loss. A study by Swift and colleagues showed that maintaining muscle strength via resistance training during HLS can also mitigate bone loss^48^. Another study in humans undergoing 17 weeks of horizontal bed rest found that resistance exercise attenuated the loss in gastrocnemius and soleus muscle mass and BMD^49^. Further, there are studies examining the effect of muscle stimulation on both muscle atrophy and bone loss. Lam and Qin reported that muscle stimulation at a mid-frequency (50 Hz) improved bone volume and structure during a 4 week HLS period^12^. Together, these data suggest that maintenance of muscle mass would attenuate unloading-induced bone loss. However, our data suggest that maintaining muscle mass without supplemental measures (i.e. exercise, dynamic muscle stimulation) does not attenuate bone loss in response to unloading or RANKL-induced bone loss. This interpretation suggests a temporal disconnect between muscle loss and bone loss in response to unloading and reinforces the need for further studies to understand the underlying mechanisms involved in the interplay between osteopenia and sarcopenia.

## ACKNOWLEDGMENTS

We thank Anne Pruznak and Gregory Lewis for their excellent technical assistance. This work was supported by the National Space Biomedical Research Institute (grant MA02802); and the National Institute on Alcohol Abuse and Alcoholism (grant AA11290).

